# A subclass of the IS*1202* family of bacterial insertion sequences targets XerCD recombination sites

**DOI:** 10.1101/2022.12.17.520857

**Authors:** Patricia Siguier, Philippe Rousseau, François Cornet, Michael Chandler

## Abstract

IS*1202*, originally isolated from *Streptococcus pneumonia* in the mid-1990s had been previously tagged as an emerging IS family in ISfinder. While searching for plasmid-associated Xer recombinase recombination sites (*xrs*) in *Acinetobacter baumannii*, we observed that some insertion sequences related to IS*1202* were repeatedly found abutting these sites in a number of plasmids. The plasmids often carried repeated *xrs* thought to form a new type of mobile genetic element (MGE) which uses the chromosomally-encoded XerCD recombinase for mobility. The MGE (*xrs* cassette) consist of *xrs* flanking one or a small number of genes often including different clinically important carbapenemase-encoding *bla*-OXA. The IS*1202-* related IS are inserted with their left, transposase proximal extremity, IRL, five base pairs from *xrs* and include a characteristic 5bp flanking target duplication. Further searches revealed that many different plasmid- and chromosome-borne *xrs* can be targeted and that IS*1202*-*xrs* combinations are not limited to *Acinetobacter baumannii* but occur in other bacteria.

In addition to 28 IS*1202* group ISs in ISfinder and a number which had been subsequently submitted, we undertook a survey of the NCBI (February 2020) and identified 138 additional IS*1202*-related IS. These could be divided into 3 principal subgroups based on their transposase sequences and on the length of the DR generated on insertion: subgroup IS*1202* 27-28bp DR); IS*Tde1* (15-17bp); and IS*Aba32* (5-6bp). Members of each group which lacked DR were also found. But since other examples of most of these were subsequently identified having DR, those lacking DR may have been generated by intra-replicon recombination. Only members of the group which generate 5bp DR were found to target *xrs*. These were not only identified in plasmids but also occurred at some individual *xrs* sites, *dif*, located at the chromosome replication terminus and involved in post-replication chromosome segregation. Further analysis showed the presence of subgroup-specific indels in their transposases which may be responsible for the differences in their behavior.

We propose that this collection of IS be classed as a new insertion sequence family: the IS*1202* family composed of at three subfamilies, only one of which specifically targets plasmid-borne *xrs*. We discuss the implications of *xrs* targeting for gene mobility.

## Introduction

The insertion sequence (IS) IS*1202* was initially identified in *Streptococcus pneumonia* in 1994 (1). It is 1,747 bp long, bordered by 23 bp imperfect inverted repeat sequences, contains a single open reading frame sufficient to encode a 54.4-kDa polypeptide and is flanked by a 27-bp direct target repeat sequence (DR). IS*1202* was not related to any of the known IS elements and was classified in ISfinder as an emerging IS family (ISNCY – not classified yet) (2) (https://tncentral.ncc.unesp.br/TnPedia/index.php/IS_Families).

A small number of IS*1202*-related IS were identified abutting Xer Recombination Sites (*xrs*) in bacterial plasmids. *xrs* are specific recombination sites found on chromosomes and plasmids and acted on by the XerC and XerD recombinases (3). They are composed of two conserved 11 bp flanking sequences which are differentially recognized and bound by XerC and XerD to form a heteromeric complex which includes two recombining *xrs* (3,4). XerC and XerD binding sequences are separated by a poorly conserved 6 to 8 bp sequence, called the central region, at which strand transfer occurs during recombination. XerCD recombine chromosome- and plasmid-borne *xrs* to resolve dimeric forms due to recombination between circular sister replicons. Other *xrs* are used to integrate bacteriophages or genomic islands into chromosomes. Lastly, numerous *xrs* have been found flanking mobile genes in plasmids, thus inferred involved in their mobility (5–8). This has been repeatedly found in plasmids of *Acinetobacter baumannii* often carrying repeated *xrs* (called p*dif* in these cases, because of their homology to the chromosomal *xrs, dif*) arranged in modules in which they flank one or a small number of genes, often including different clinically important carbapenemase-encoding *bla*-OXA genes (5–8).

During plasmid-borne *xrs* annotation and characterization, we identified a large number of IS insertions abutting these sites in replicons, both chromosomes and plasmids, of many bacterial genera and species including, but not exclusively, *Acinetobacter baumannii, Klebsiella pneumoniae* and *Burkholderia cenocepacia* (supplementary Table 1). Interestingly, several studies had identified certain IS*1202*-related IS abutting *xrs* in plasmids of *Acinetobacter baumannii* (6,7,9). These occur at a distance of 5 bp, the length of the direct target repeat generated by insertion of these IS.

We have now identified 166 members of this emerging IS family, IS*1202*, which we analyze below. These fall into a number of subgroups defined by their transposase (Tpase) signatures and by the length of the direct target repeats (DR) that they generate upon insertion. We have named each of the three principal major subgroups after one of their members as: IS*1202*, IS*Tde1* and IS*Aba32. xrs* targeting appears to occur with members of only one of these sub-groups, IS*Aba32*, which generate 5-6 bp DR. We significantly extend the analysis of *xrs* targeting by identifying 125 examples of IS insertions abutting *xrs*: these insertions occur not only in *Acinetobacter* plasmids but can be identified in plasmids of other bacterial genera such as *Klebsiella, Burholderia, Bradyrhizobium* and *Serratia* (supplementary Table 1). They involve both different insertions of the same IS as well as different IS of the same subgroup and are found not only in *pDif* cassettes but also occur abutting resident single chromosomal *dif* sites as well as additional single *xrs* sites in the chromosomes of these and other genera. Moreover, several examples of *xrs*-associated tandem IS insertions have been identified.

## Results

### Identification of IS*1202* family members

Approach, numbers, distribution.

At the beginning of this study (February 2020), the ISfinder database included 21 sequences belonging to the emerging IS*1202* group (previously ISNCY; (10)). The direct repeats generated by their insertion varied in size from 0 to 28 nucleotides. Seven additional examples were subsequently submitted to ISfinder by: Moran, 2021; Harmer and Hamidian, 2021; Feng 2021; and Siguier 2020.

We further enriched the library by identifying 138 IS*1202*-related IS from public databases at NCBI using reiterative BLAST approaches as described previously (similar to that used in (11)) with the primary transposase sequence of representative elements used as a query in a BLASTP (12). The collection of IS clearly forms a coherent family, the IS*1202* family, based on their transposase sequence.

Comparison of the 166 members showed that they range from 1,320 to 1,990 bp in length except for 5 examples (IS*Kpn21*, IS*Kpn65*, IS*Kpn63*, IS*Shal2* and IS*Rel10* which include an unrelated passenger gene annotated as “hypothetical protein”) with a single Tpase *orf* of between 400 and 500 amino acids in a single reading frame (Fig. 1A).

**Figure 1.**
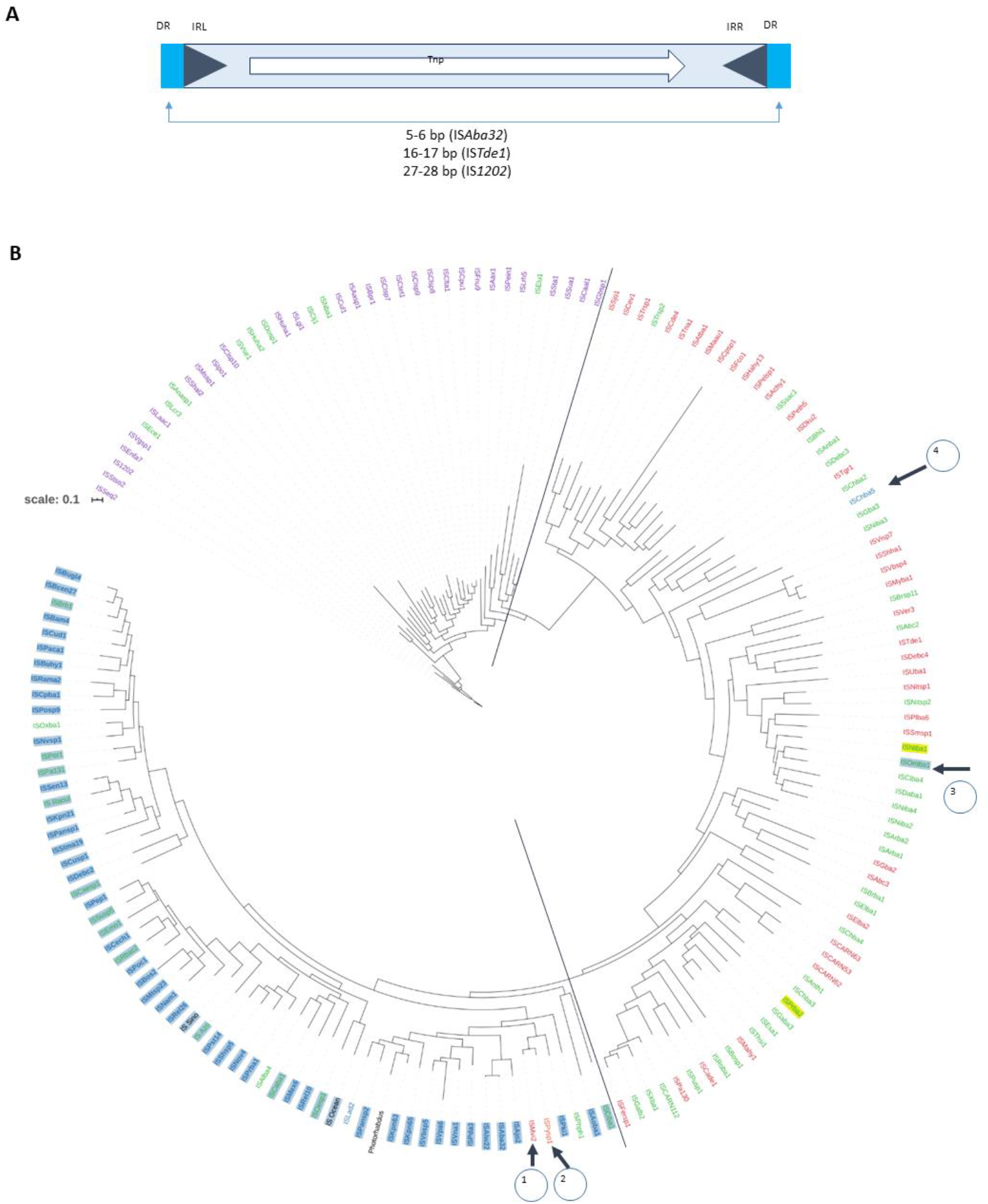
General Structure of members of the IS*1202* family and their Phylogenetic Tree. **A)** The IS is shown as a blue box with blue triangles indicating the left and right inverted repeats, IRL and IRR. The transposase, Tnp, open reading frame is shown within the box as a white arrow to indicate the direction of expression. Flanking Direct target repeat sequences, DR, are indicated by light blue boxes. The DR length of the three IS1202 sub groups is shown below. **B)** The tree is based on transposase amino acid sequences of 166 IS. The IS names are colored coded according to the length of the DRs they generate: lavender, very long DR, 27-28bp; red, long DR, 16 - 17 bp; blue, short DR, 5-6bp. Names marked in green are IS which do not have DR. Those boxed in blue are IS which are located next to an xrs site. There is a clear correlation between length of IR and xrs. The black arrows indicate individual IS which vary slightly from the overall pattern

For each IS, we annotated at least one (and frequently several) insertion sites corresponding to insertions in different loci. This generated a library of 229 insertions which include the flanking 100 bp (Supplementary Table 1).

### Phylogenetic Analysis and the identification of subgroups

To classify these IS sequences further, we analyzed their characteristics: insertion sites, transposase sequence, their IRs and the length of the DRs which they generate.

ISs of IS*1202* family generate three types of direct target repeats with different lengths: 5-6 bp (61 examples), 16-17 bp (67 examples) and 27-28 bp (38 examples) (Fig.1A). In the case of a few IS, no direct repeats were present. However, in many cases, other copies of the IS did exhibit DRs. The absence of DR in these cases could therefore simply be the result of intra-replicon recombination between two resident IS copies, leading to the separation of the flanking DR sequences or simply result from genetic drift.

A phylogenetic tree based on the transposase amino acid sequences generated using MAFT (Materials and Methods) and viewed with iToL (Interactive Tree Of Life; https://itol.embl.de/) is shown in Fig. 1B. The IS are clustered into three principal groups: IS*Aba32*, IS*Tde1* and IS1*202*. Interestingly these three groups are also distinguished by the length of the DR they generate: IS*Aba32* (5-6 bp), IS*Tde1* (16-17 bp) and IS1*202* (27-28 bp). Several carry apparently unrelated passenger genes. These are not restricted to a single subgroup: for example, whereas IS*Kpn21*, IS*Kpn65*, IS*Kpn63*, and IS*Rel10* all belong to the IS*Aba32* group, IS*Shal2* (29 bp DR) belongs to the IS*1202* subgroup.

The distribution of these groups is also quite different (Supplementary Table 1): members of the IS*Aba32* subgroup can be found in both the plasmids and chromosomes as well as in unassembled shotgun sequences of mainly proteobacteria of the γ (Acinetobacter) and β (Burkholderia) and some α proteobacteria. The majority of IS*Tde1* subgroup members were identified in whole shotgun sequences, in a number of chromosomes but in only one plasmid. They are more disperse and can be found in γ and β proteobacteria, Firmicutes, Armatimadetes, Deltaproteobacteria, Nitrospira, Gemmatimonadetes, Elusimicrobia, Chloroflexi, Synergistetes, Acidobacteria and Atribacterota. While the IS*1202* subgroup is found in whole shotgun sequences and also in assembled chromosomes. We have not yet identified plasmid copies of this subgroup. They are found in Firmicutes, Tenericutes and one Spirochaete (Supplementary Table 1).

### Transposase Signatures

To identify transposase domains, we performed a domain search with COG and HMMER/PFAM and a *de novo* search with MEME (Materials and Methods). This revealed two major domains: an N-terminal helix-turn-helix (HTH) DNA-binding domain and a DDE-type RNase fold catalytic domain (Fig 2 and Supplementary figure 1). All three subgroups also included a non-conserved C-terminal region (Figs. 2B).

**Figure 2.**
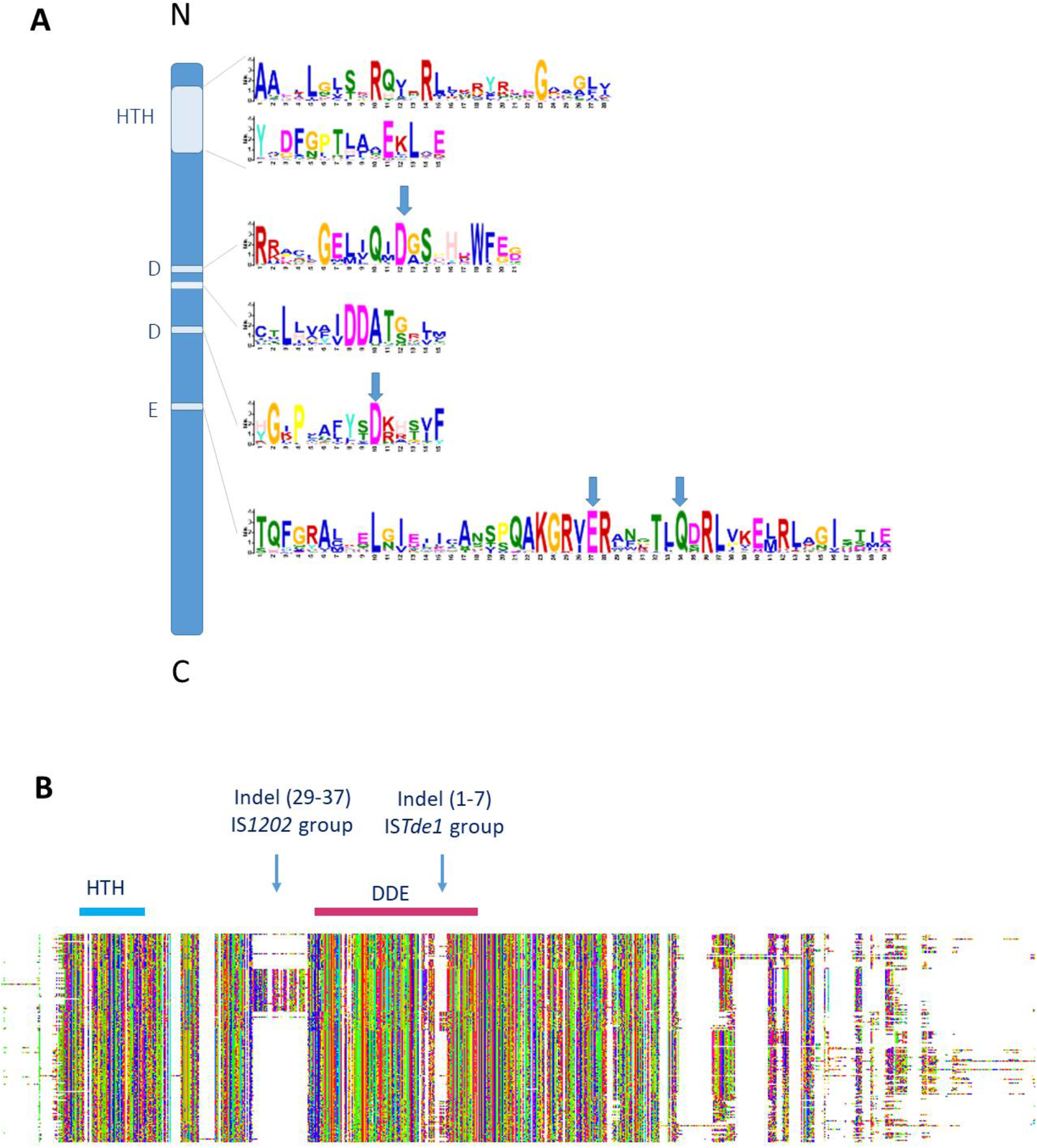
Schema of the transposase domain with the 6 conserved motifs. A) The six conserved motifs revealed by MEME are shown as a function of their position along the transposase. The N-terminal and C-terminal ends of the protein are indicated as are the helix-turn-helix motif (HTH) and the DDE triad and the two additional conserved D residues. The conserved residues of the DDE motif are indicated by vertical blue arrows. B) **Transposase Alignment using MAFFT and visualized with MSAviewer**. XXX transposase sequences are included in the alignment. The position of the indels is indicated as well as that of the HTH ans DDE motif together with the number of IS containing each indel in parentheses. Note that the indel found in the ISTde1 group occurs within the DDE motif. The non-conserved C-terminal and of these proteins is clearly indicated. The colors represent the level of conservation

Closer inspection of these domains (Fig. 2A) showed the presence of two highly conserved additional Aspartic acid (DD) residues between the two D of the DDE domain. They also include a glutamine (Q) seven residues C-terminal to the conserved E (Glutamic acid) instead of the characteristic K/R (Lysine/Arginine) (2,10). The motifs surrounding the DDE triad are retained by each of the subgroups individually (Supplementary figure 2).

The MAFFT alignment with the transposases retained for the study (Materials and Methods; Fig. 2B) revealed two prominent group-specific indels: one of about 30 amino acids just before the catalytic domain (38 examples from the IS*1202* subgroup), and second smaller indel of 5-10 amino acids (43 from the IS*Tde1* subgroup) between the second D and the E of DDE domain. There was significant amino acid conservation in the larger indel particularly at the N-terminal end (Supplementary figure 3).

To obtain some indication of the impact of these indels might have on transposase organization, we used Alpha fold (13) to generate potential structures (Fig. 3). Although it must be emphasized that these are only models, they reveal the N-terminal HTH (Helix-Turn-Helix) domain, presumably involved in IR binding, separated from the catalytic domain carrying the catalytic site by a poorly defined segment (blue arrows). The variable C-terminal segment, predicted to be α-helical is also poorly defined (orange arrows) as a region of low or very low predictive confidence. The positions of the indels in IS*1202* and IS*Tde1* transposases are indicated by red boxes and their relative positions on the IS*Aba32* scaffold are indicated by green arrows. Note that in the case of IS*Tde1*, the insertion splits the DDE motif. In all three cases, the N-ter HTH appears to be separated by a region of low predictive confidence.

**Figure 3.**
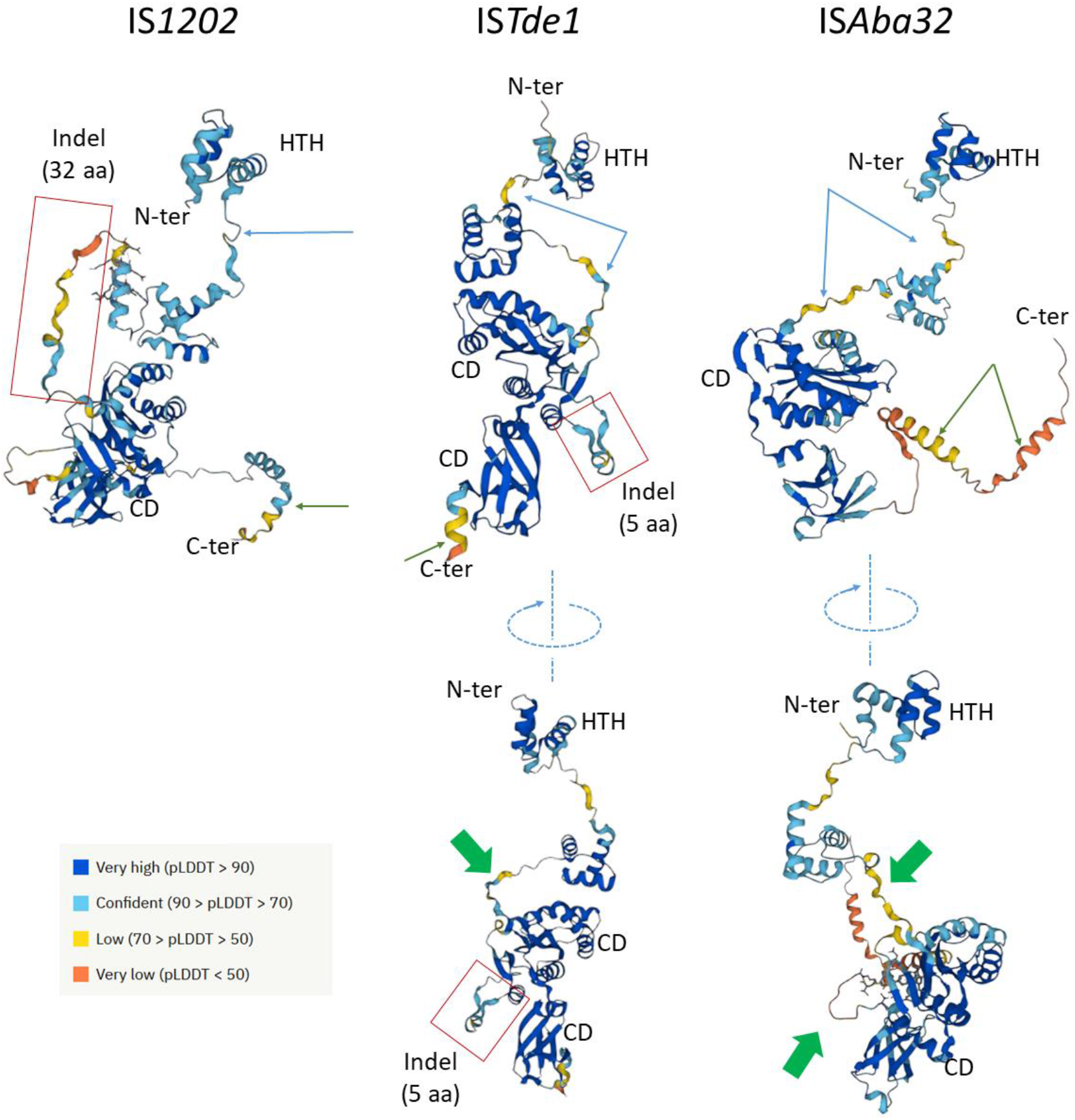
Results of Alphafold modeling. *The figure shows the predicted structure of a representative example, IS1202, ISTde1 and ISAba32, of each of the three IS1202 subgroups using the NCBI accession number for each. The N- and C-terminal ends are shown where visibleas are the catalytic domains containing the RNase fold and the DDE motif (CD) and the probable N-terminal HTH DNA binding domain. The color scheme shows the degree of certainty of the different regions of the model: dark blue, high; light blue confident; yellow, low; orange very low. Red boxes indicate the position of the indels. Their positions on the scaffold of ISAba32 which include neither is indicated by a green arrow*.

The transposases appear to be distantly related to IS*481* and IS*3* family transposases particularly in their DDE domains (e.g. IS*1202* has 39% amino acid similarity to IS*Pfr5* of the IS*481* family) (10).

### The Terminal Inverted Repeats

We also analyzed the terminal inverted repeats of the collection. For each IS sub group, we aligned the left and right ends (Fig. 4). Like the IRs of most IS (see: https://tncentral.ncc.unesp.br/TnPedia/index.php/General_Information/IS_Organization), IS*1202* family IRs carry two well conserved domains: a terminal domain of three base pairs, which is recognized for cleavage, and an internal region which generally serves as a DNA recognition sequence for transposase binding. The terminal domain of both IRL and IRR of two subgroups (IS*Aba32* and IS*Tde1*) begins with 5’-TGT-3’ (as do those of the IS*3* and IS*481* families (10)) while those of the third subgroup, IS*1202*, are less conserved: IRR retains the conserved TGT, but the left end is less conserved (5’-Ta/gT-3’). All three carry the second conserved region around position 20 at both ends. This is somewhat more extensive for the IS*1202* group than for the other two groups. Not only does the IS*Aba32* group carry a third relatively well conserved region further into the IR, but exhibits a completely conserved C residue at position 9 on IRL.

**Figure 4.**
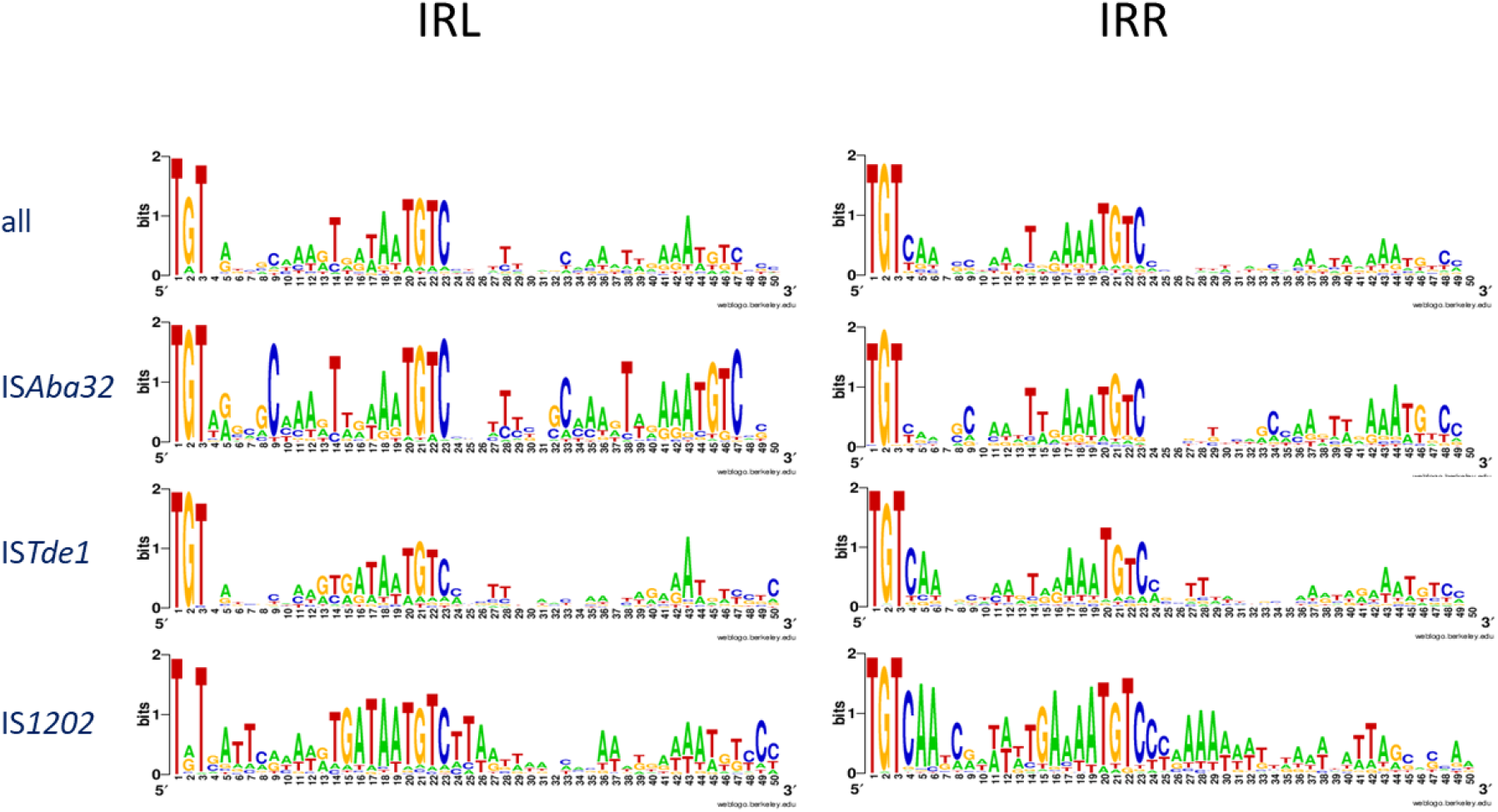
Alignment of IRL and IRR. The sequences of IS ends were aligned using WebLogo. They are defined by the direction of transcription of the transposase gene. IRL, by definition, is located on the 5′ side of the transposase orf. Top: Alignment of 213 left (IRL) and right (IRR) ends including all three IS1202 subgroups. Alignment of those of the individual subgroups are shown below: ISAba32, n=96; ISTde1, n=70; IS1202, n=47.

Thus, in addition to the DR length and specific indels, each of the IS*1202* family subgroups can also be distinguished by their terminal IRs.

### *xrs* Targeting

Perhaps the most striking characteristic of the IS*Aba32* subgroup is its insertion specificity. In the framework on our survey of *xrs* sites present in *Acinetobacter* plasmids, we identified a large number of IS insertions abutting these sites (Supplementary Table 1; Fig. 1B, Fig. 5). We subsequently observed these types of insertion in *Klebsiella pneumoniae* and *Burkholderia cenocepacia* among a large number of bacterial species and genera (supplementary Table 1). The IS is invariably inserted at a distance of 5 to 6 bp from the *xrs*, corresponding to the length of the direct repeat (DR) generated by IS insertion. In each case, the IS insertion was oriented: the left end of, IRL, was always to the right side of the *xrs*, the XerC arm (Fig. 5A). We found both full length and partial copies of IS*Aba32*, in several plasmids (Supplementary Table 1) and in proximity to different *xrs*. In each partial copy, IRL (but not IRR) was conserved and the distance between the *xrs* and the partial IS remained 5 or 6 bp. The same IS could also insert next to different *xrs* sites: for example, 3 copies of IS*Ajo2* are inserted next to 3 different *xrs* (Fig. 5Bi) (as judged by the variation in their central regions in *A. baumannii* plasmid pAF-401 (Fig. 5A and 5Bi). This type of targeting has previously been observed only in a limited number of cases in *Acinetobacter* strains (6,7,9).

**Figure 5.**
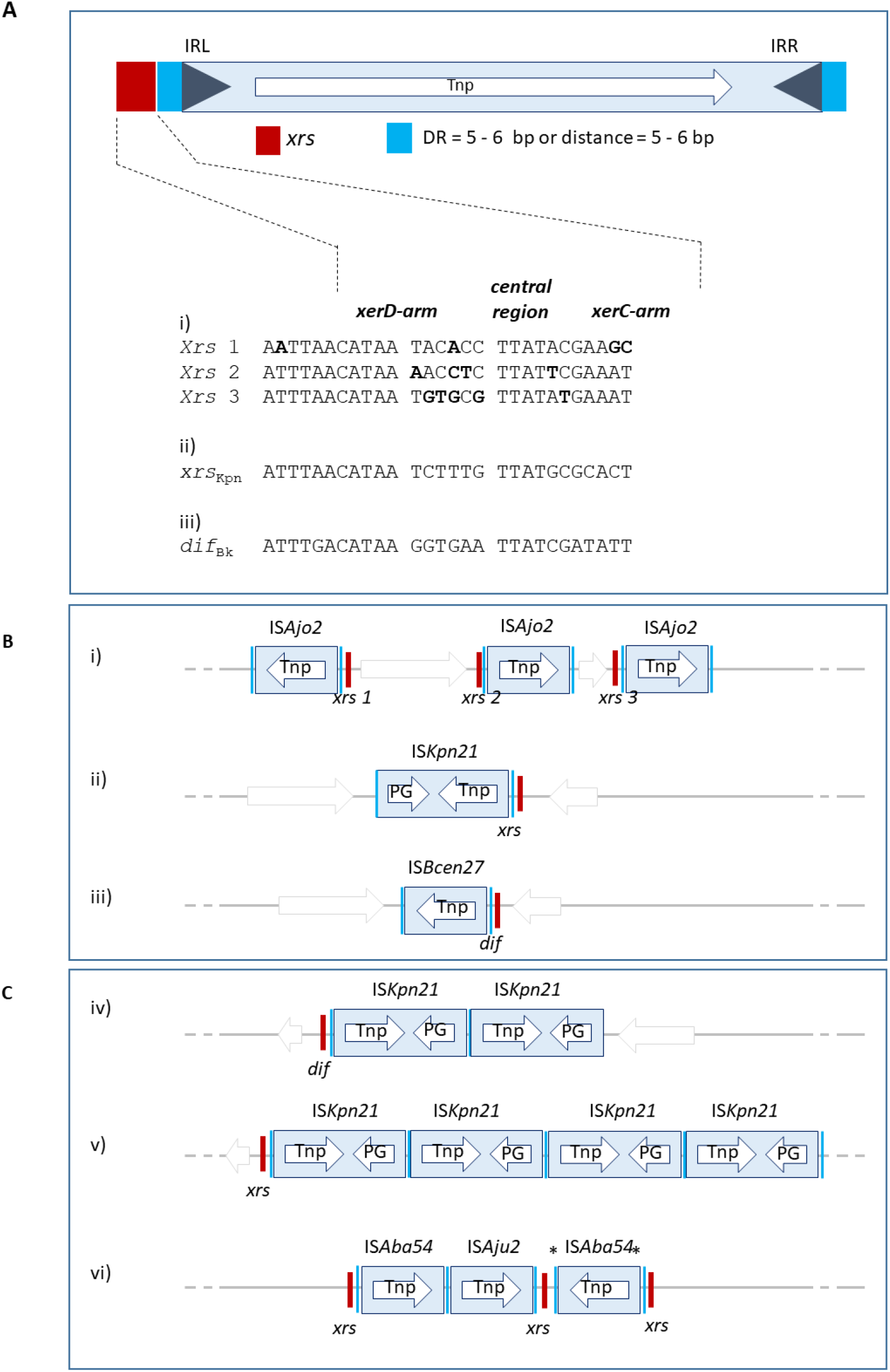
Targeting of members of the IS*Aba32* group to xrs sites. **A) Position of the IS with respect to an xrs site**. The IS is shown as a blue box with blue triangles indicating the left and right inverted repeats, IRL and IRR. The transposase, Tnp, open reading frame is shown within the box as a white arrow to indicate the direction of expression. Flanking Direct target repeat sequences, DR, are indicated by light blue boxes. The xrs site is shown to the left with the xerD arm shown in red and the xerC arm in salmon. Below: xrs site sequences observed in examples Bi,ii and iii) with the xerC, central region and xerD arms indicated. **B) Examples of different ISAba32 members in plasmid and chromosomes**. i) Acinetobacter baumannii AF-401 plasmid pAF-401 (NZ_CP018255) carrying three copies of ISAjo2 at three different xrs sites. The xrs site sequences are shown above. ii) Klebsiella pneumoniae CAV1193 plasmid pCAV1193-258 (CP013323). iii) Burkholderia cenocepacia MC0-3 chromosome 1 (CP000958). **C) Examples of targeting by several ISAba32 members near the same xrs**. iv) Klebsiella pneumoniae ARLG-3226 (CP067826): two copies of ISKpn21 after the dif site of the chromosome. v) Klebsiella pneumoniae isolate 307 genome assembly, plasmid: P1 (OX030709): four copies of ISKpn21 near only one xrs. vi) Acinetobacter junii strain ZM06 plasmid unnamed (CP077416): targeting of one xrs by two different ISAba32 members and insertions of two members of ISAba32 group after one xrs.

We next asked whether other (all) members of this family target *xrs* by searching for the presence of an *xrs* site neighboring each IS in our library. Strikingly, only ISs of the IS*Aba32* subgroup were inserted next to *xrs* (represented by blue boxes in Fig.1B, supplementary Table 1). Members of the other two IS*1202* subgroups did not show this targeting behavior. Of the 166 members of our IS*1202* library, 61 were inserted at 5-6 bp from an *xrs*. All belonged to the IS*Aba32* subgroup including at least one, IS*Kpn21*, which carries a passenger gene (Fig. 5Bii). The insertion was always oriented: with the left IS end to the right side of the *xrs* XerC arm (Fig. 5A). Of these, 57 had a DR of 5-6 bp and 4 had inserted at the correct distance but did not have an identified DR. It should be noted that, for convenience, *xrs* are shown inverted in certain figures (*e*.*g*. see Fig. 6A below).

**Figure 6.**
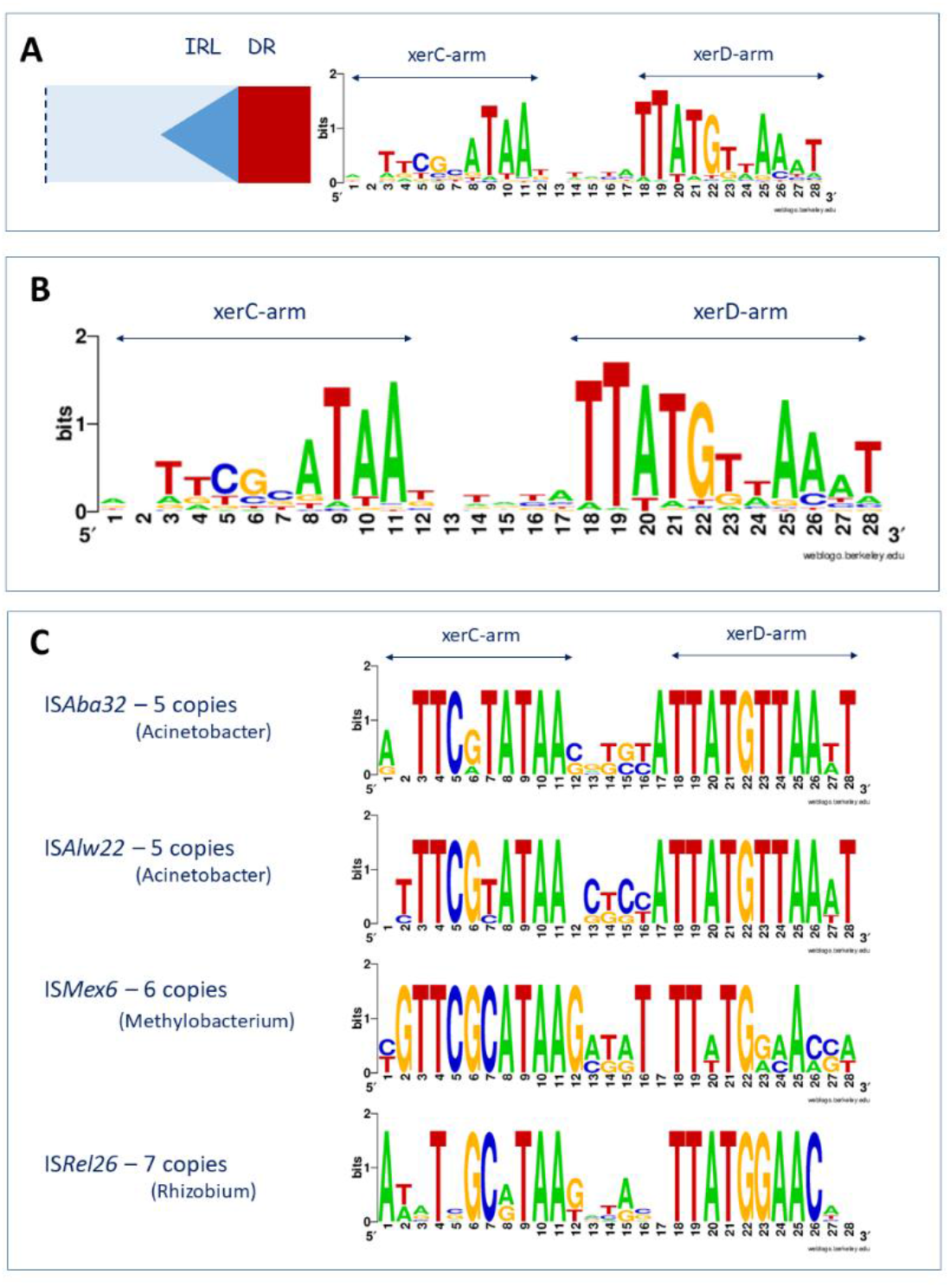
Detailed view of *xrs* sites. **A) Orientation of xrs sequences shown in Weblogo format with respect to ISAba32 subgroup members**. **B) Alignment of 78 individual xrs sequences in WebLogo format**. This indicates that the sequences, and particularly the central regions, are not conserved demonstrating that different ISAba32 subgroup members can target a number of different xrs sites. **C) Alignment of xrs sequences found next to the same ISAba32 subgroup member**. ISAba32, Acinetobacter, n=5; ISAlw22, Acinetobacter, n=5; ISMex6, Methylobacterium, n=6; ISRel26, Rhizobium, n=7. Neither the central regions nor the XerC and XerD binding sites are conserved.

To determine whether the IS*Aba32* subgroup targets different *xrs*, the repeated copies of insertions into the same *xrs* were removed from the 166 insertion and the remaining 78 *xrs* sequences were aligned to obtain the WebLogo of Fig. 6B. The targeted *xrs*, particularly their central regions, are not conserved. This demonstrates that different IS*Aba32* subgroup members can target a number of different *xrs* sites.

To determine whether a particular IS recognizes a particular *xrs* site, we aligned the *xrs* site present next to different copies of the same IS (Fig. 6C). The central regions are not conserved, and the XerC and XerD binding sites are not identical. It is therefore not the nucleotide sequence alone that is recognized by the IS.

We also identified a number of chromosomally located insertion sites of IS*Aba32* subgroup members. A selection on these are presented in Fig. 7 on a cumulative GC skew plot of the chromosome to indicate the positions of origin and terminus of replication (14). Most of these sites are chromosomal *dif* sites indicating that insertion can target *xrs* acting in chromosome dimer resolution. Examples include: IS*Bcen27* present in two of the *B. cenocepacia* MC0-3 chromosomes and also at two positions in one of the *B. cenocepacia* PC184 Mulks chromosomes; IS*Kpn21* which occurs in several *Klebsiella* and *Serratia* strains; as well as other IS in *Bradyrhizobium* and in a number of chromosomes of various *Acinetobacteria* (Supplementary Table 1).

**Figure 7.**
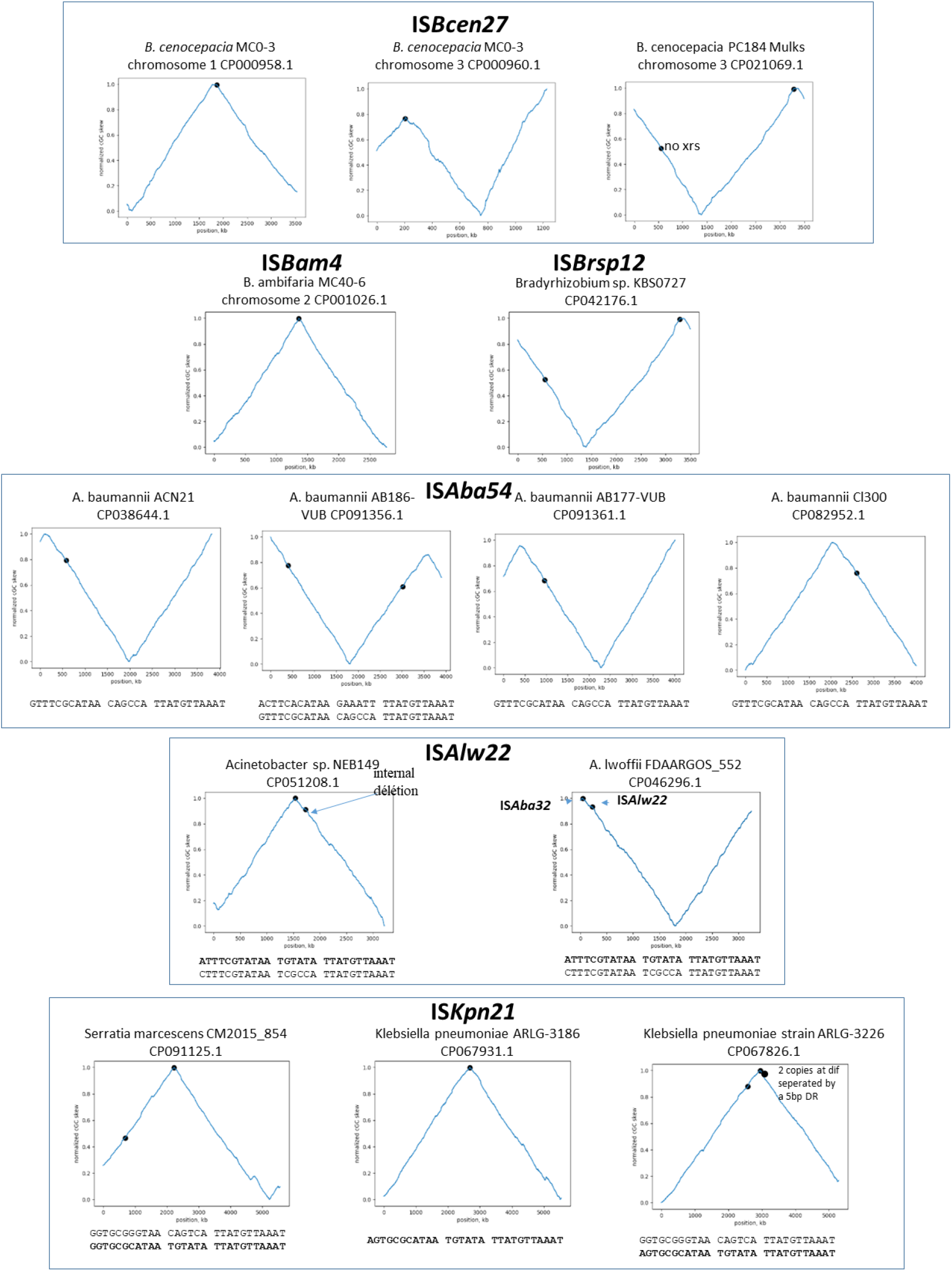
Position of IS neighboring *xrs* sites on bacterial chromosomes. Examples of chromosomes shown as cumulative GC skew maps. In this view, the terminus of replication occurs at the peak and the origin at the trough. The position of the IS are shown as blue dots next to xrs (principally dif) sites. The chromosome examples are grouped according to the particular IS involved (indicated in each box with the name of the host and its chromosome number if there are multiple chromosomes). In the case of ISKpn21 (bottom left, two tandem copies were identified.

Finally, a number of *xrs* sites are abutted by several IS (fig 5Civ-iv) indicating the *xrs* serves as a target for successive IS insertions. This is the case for *K. pneumonia* ARLG-3226 which carries two tandem IS*Kpn21* copies at *dif* (although the distal IS*Kpn21* copy is missing a flanking DR) (fig. 5Civ) in addition to a third copy at an *xrs* site at some distance. A second example is *Klebsiella pneumoniae* isolate 307 plasmid P1 (OX030709) (Fig. 5Cv) which carries 4 IS*Kpn21* copies each separated by the same 5bp DR. If these insertions were targeted to the *xrs* site, it implies that the *xrs* proximal IS copy was the last to arrive. The third example from an unnamed plasmid in *Acinetobacter junii* strain ZM06 (CP077416) is more complex (Fig. 5Cvi): there is one copy each of IS*Aba54* and IS*Aju2* separated by the 5bp DR. There is also a second copy of IS*AbA54* inserted next to a second *xrs* with a 5bp DR whose sequence is different from the others (marked *). Note that there is a third non-contiguous *xrs* copy in this plasmid, which has not been involved in IS insertion. This example shows that different IS can target the same *xrs* and the structure again implies that the *xrs*-proximal IS arrived last.

## Discussion

We have expanded the group of IS including IS*1202*, previously part of the IS*NCY* (Not Classified Yet) section of ISfinder (2,10) to include 138 related IS. This was achieved using a reiterative BLAST protocol (11). The collection forms a coherent IS group which we propose to call the IS*1202* family.

Members of this family have impacted some important properties of their hosts. For example, it was observed (15) that an IS*1202*-related IS was implicated in the deletion of the capsular polysaccharide locus (*cps*), the major known *Streptococcus pneumoniae* virulence factor, important for Streptococcal survival in the blood and strongly associated with antiphagocytic activity (16). While Chamoun et al (17) showed that insertion of IS*Ajo2* into the *lpxA* gene, involved in lipid A biosynthesis, can result in colistin dependence which, they suggest, possibly leads to colistin resistance.

Transposase alignment and phylogenetic analysis of these IS*1202* family members identified 3 major subgroups: IS*Aba32*, IS*Tde1* and IS*1202* (Fig. 1B). These all exhibited an N-terminal HTH domain and a catalytic domain which includes a pair of conserved D residues between the first and second D of the DDE triad (Fig. 2) and a particular Q residue downstream from the E replacing a R/K residue, a pattern which had been noted previously (2,10). The subgroups could also be distinguished by the presence of group-specific indels (Fig. 2C) and were also clustered according to the length of the direct target repeats they generate on insertion (5-6bp: IS*Aba32*; 15-17bp: IS*Tde1*; and 27-28bp: IS*1202*) (Fig. 1B). A number of these did not exhibit DRs. This might due to recombination between two identical copies of the IS which would result in distribution of the DRs between the two recombining partners.

The indels occur between projected structural domains of the proteins as indicated by structural models derived using the α-fold software (Fig. 3). These indels are associated with mechanistic changes in transposition strictly associated with the behavior of the IS*1202* subgroup in which they are found *viz*: loss of *xrs* targeting and increase in the length of the associated DR.

### The IS*Aba32* group and sequence-specific targeting of *xrs*

The original ISfinder library (February 2020) contained 21 examples of IS*1202*-related IS. Eight of these generated 5 bp DR and were located next to an *xrs* site. Seven additional IS described in references (6,7,9), all restricted to *Acinetobacter* species, were also observed to have inserted near a plasmid-associated *xrs* site. Here we have greatly expanded the number of examples which are associated with *xrs* sites to include a significant number of different bacterial genera and species. We found examples integrated into plasmid-associated ‘p*Dif* cassettes’ which consisting of gene-carrying DNA fragments flanked by inversely-repeated *xsr* and appear mobile, both within and between species (18,19), although the way they move is currently unclear. We also identified examples inserted next to individual chromosomal *dif* sites (Supplementary Table 1). Even those few individual IS examples identified which did not obviously occur next to such sites proved to have identical sister copies elsewhere which were associated *xrs*.

It is also clear that a given IS can target different *xrs* sites (Fig. 5Bi) and that one *xrs* site can act as a target for multiple insertions of identical and different members (Fig. 5Civ-vi) of the IS*Aba32* subgroup.

The mechanism of IS*Aba32* subgroup targeting of *xrs* sequences is at present a matter of speculation. It could be the result of direct transposase interactions with the XerC and or XerD proteins themselves or to a direct recognition of *xrs* architecture. Neither is it clear why insertion is directional i.e. that it is always IRL which abuts the *xrs* XerC arm: clearly IRR is less well conserved than IRL, particularly in the internal region (Fig. 4).

One possible advantage of targeted insertion to *xrs* sites is that insertion could increase expression of a downstream gene either by forming a hybrid promoter (20)(https://tncentral.ncc.unesp.br/TnPedia/index.php/General_Information/IS_and_Gene_Expression) or by providing a mobile promoter (e.g. IS*Ecp1*; (21) https://tncentral.ncc.unesp.br/TnPedia/index.php/IS_Families/IS1380_family). The IS orientation with respect to neighboring *orfs* is, however, often not compatible with this. Another possibility is that they are a safe haven as in the case of insertion of the Tn*7* transposon directed by an attTn*7* sequence downstream from the highly conserved *glmS* gene (see (22)). It is important to point out that some *xrs* are flanked on their XerC-side by specific regions (called ‘accessory regions’) containing binding sites for various accessory proteins which serve an architectural role and control XerCD-mediated recombination (23–25). As a particular DNA structure, accessory regions might be targeted by the IS. We do not know at present whether the targeted *xrs* possess such flanking elements that have been described only in enterobacteria so far. In addition, targeting may inactivate possible accessory region-mediated control, damaging dimer resolution at these sites. However, the fact that ISs target plasmid-borne *pDif*-cassettes and chromosome *dif* sites, neither of which is predicted to use this kind of control, argues against this possibility.

However, it is important to point out that *xrs* sites are sometimes flanked on one side by sites for various accessory factors which serve an architectural role in defining the shape of the nucleoprotein recombination complex (23–25). Alternatively, the IS could act as accessory sites themselves, facilitating formation of the appropriate topology required for recombination. That the targeted *xrs* sites in cassettes are probably active is supported by the observation that they are recognized by *Acinetobacter baumannii* XerC and XerD *in vitro* (Blanchais et al., in preparation).

### Transposition mechanisms

Little is known concerning the transposition mechanism of this IS family. However, there are two reports in which circular IS copies have been identified. Hudson et al (26) identified circular copies of IS*Kpn21*, a member of the IS*Aba32* subgroup, during analysis the antibiotic resistance genes of a clinical *Klebsiella pneumoniae* ATCC BAA-2146 carbapenem resistant isolate carrying the metallo β-lactamase, NDM-1. MiSeq reads were found where IS*Kpn21* ends were linked, and separated by 5-bp direct repeats. PCR was used to rule out that this was due to tandem IS*Kpn21* copies in the host genome. This product is typical of the circular intermediates generated by a number of IS families by a mechanism called copy-out-paste-in (27) in which one IS end attacks the other, several base pairs into the flanking donor DNA. Moreover, the circle appeared to be derived from a resident plasmid copy of IS*Kpn21* which has different 5bp target flanks (possibly resulting from inter IS recombination): Only the left end flank was observed between the IS ends in the circle, suggesting that the right end preferentially attacks the left during circularization (rather than the left end attacking the right as stated by these authors)(see (28,29)). The second example was described by Nielsen et al (30) in a study of *Sphingobium herbicidovorans* MH in which they identified a circular form of an IS related to IS*1202* in addition to IS*3*, IS*6* and IS*110* family circular forms with abutted left and right ends. Unfortunately, the DNA sequence of the collection of circles is not available in an assembled form and it is therefore not possible to identify the IS*1202*-related IS. It should be noted that IS which have adopted the copy-out-paste-in mechanism, there is generally an outward-facing -35 promoter element located in the right end and an inward-facing -10 element in the left end. This results in the temporary formation of a strong promoter in the circular intermediate which permits high levels of transposase expression (for review see (27)). This appears to be the case for IS*Kpn21* and may be general for the IS*Aba32* subgroup. It remains to be seen whether the copy-out-paste-in transposition pathway is a general mechanism adopted by the entire IS*1202* family.

Finally, the target specificity of the IS*Aba32* subgroup might provide useful in targeted gene insertion protocols for introduction of various genes at pre-defined chromosomal locations.

## Aknowledgements

We would like to thank members of the GeDy laboratory at the LMGM-CBI, in particular, Manuel Campos for help with the GC skew analysis and Jocelyne Perochon (CDI) for informatics support with ISfinder. This work was supported by grant ANR InXS.

## Materials and Methods

### Identifying IS1*202* Family Insertion Sequences

IS1202-related insertion sequences were identified by reiterative blast analyses (11)(Siguier et al., 1595) with the primary transposase sequence of representative elements used as a query with BLASTP (12)(Altschul et al., 1990) at NCBI. Four sequences from the ISfinder IS*1202* group, IS*1202*, IS*Aba32*, IS*CARN112*, IS*Seq2 et* IS*Pein1*, were chosen as seed sequences. Sequences were retained if the transposases were 98% similar (identity and similarity) with transposases already in the ISfinder IS*1202* group (February 2020).

The corresponding DNA together with 1000 base pairs upstream and downstream was extracted and examined manually to identify the IRs and flanking DRs. In cases where more than a single IS copy was identified, BLASTN was used to define the IS ends. Where only a single copy was found, the ends were often defined by identification of and comparison with empty sites in other replicons. IS ends were aligned using WebLogo (31).

### Identification of xrs

BLASTN using 100 bp IS-flanking sequences as query against two in-house libraries of annotated *xrs* sites composed of 182 identified in the Enterobacteriacae (Cornet unpublished data) and 118 from Acinetobacter (Siguier unpublished). To facilitate further semi-automatic *xrs* identification, the annotated *xrs* were included in a SnapGene (www.snapgene.com). In certain cases, *xrs* sites neighboring IS could be identified manually and were added to the collection.

The xrs sites were then aligned using WebLogo (31)

### Transposase Analysis

Transposase sequences were anlaysed via the Galaxy Pasteur platform (32)(https://galaxy.pasteur.fr/) and NGPhylogeny (https://ngphylogeny.fr/). They were aligned using MAFFT (33) and trimmed with TrimAl (34). A phylogenetic tree was constructed with PhyML (35) and visualized with annotations (presence and length of DR and presence of *xrs* in proximity to the IS) using iTOL (36)(https://itol.embl.de). Transposase domains were identified with CDD-NCBI (https://www.ncbi.nlm.nih.gov/Structure/cdd/cdd.shtml), PFAM/HMMER (https://www.ebi.ac.uk/Tools/hmmer/search/hmmscan) or *de novo* using MEME with searches for three or six motifs (https://meme-suite.org/meme/).

Predictive transposase structural models were obtained using Alphafold (13). These can be accessed through the following links :

IS*Aba32* (DR=5 pb) : https://alphafold.ebi.ac.uk/entry/A0A5N5XUG9;

IS*Tde1* (DR= 17 pb) : https://alphafold.ebi.ac.uk/entry/Q73JR2;

IS*1202* (DR= 27 pb) : https://alphafold.ebi.ac.uk/entry/Q54513.

### Chromosomal xrs Positions

The position of IS-occupied chromosomal *xrs* sites was visualized on a cumulative GC skew map (14) using in-house Python software (Manuel Campos, personal communication).

## Supplementary Material

**Supplemental figure 1.**
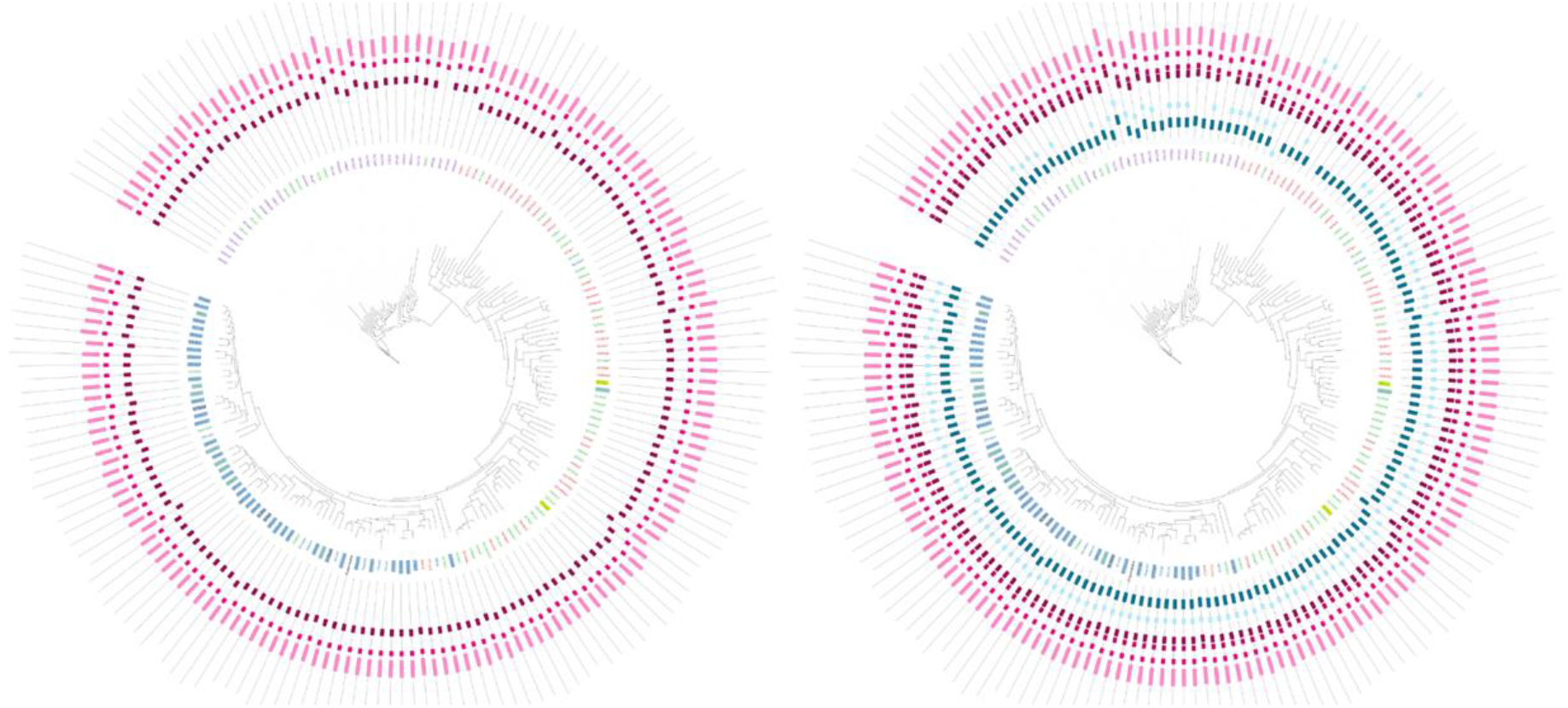
Position of the *de novo* domains identified with MEME and visualized with iTol on the phylogenetic transposase tree. The left segment is the result using a maximum of 3 domains, revealing the DDE triad (pink) and the right segment shows the results of using a maximum of 6 domains which reveals, in addition the conserved DD region within the DDE domain (pink) and the HTH domain (blue)

**Supplemental figure 2.**
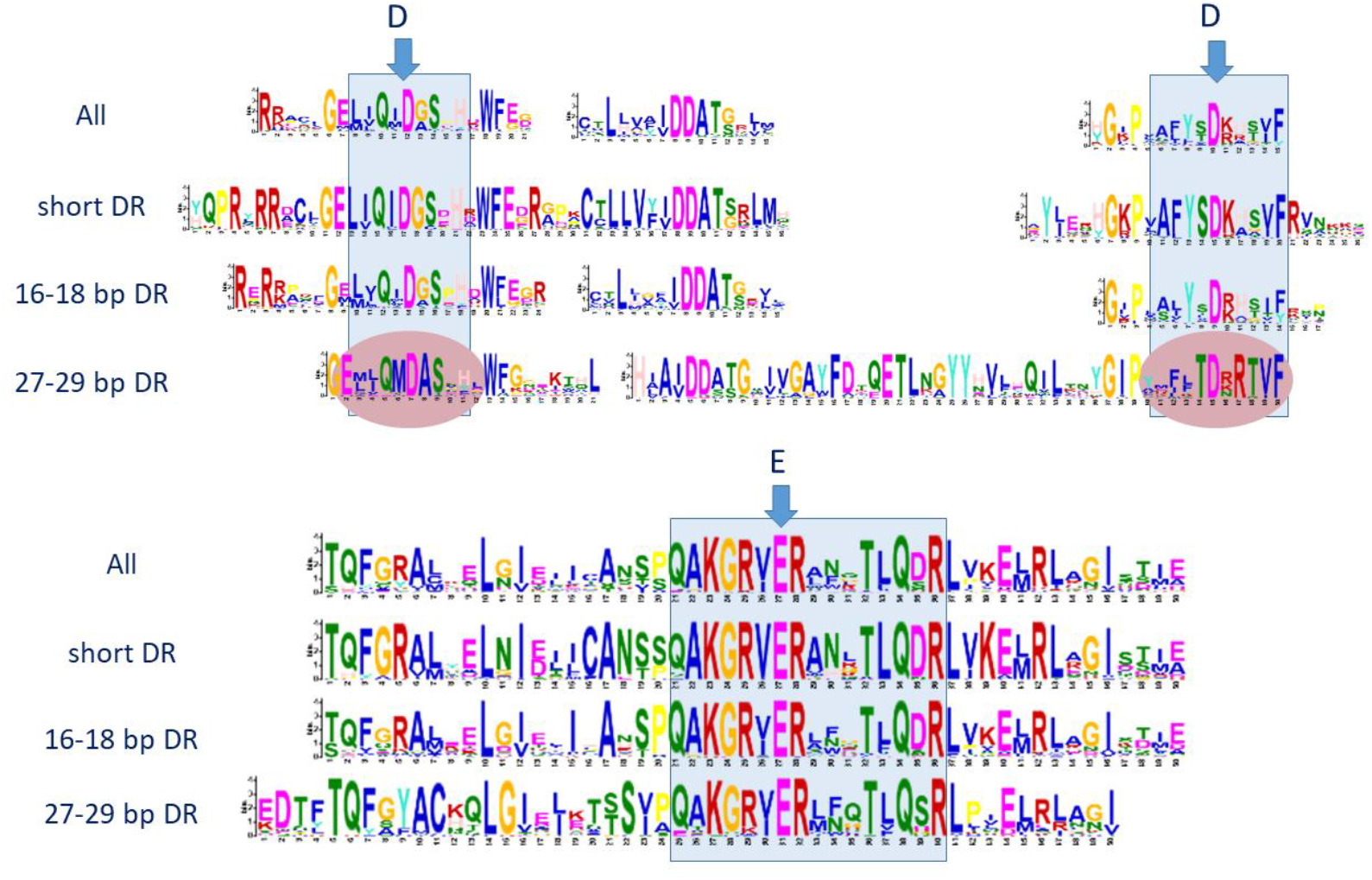
Schema of the transposase domain with the 6 conserved motifs for each individual IS*Aba32* subgroup. The DDE motif revealed by MEME are shown in blue boxes together with the conserved additional DD. The conserved residues of the DDE motif are indicated by vertical blue arrows.

**Supplemental figure 3.**
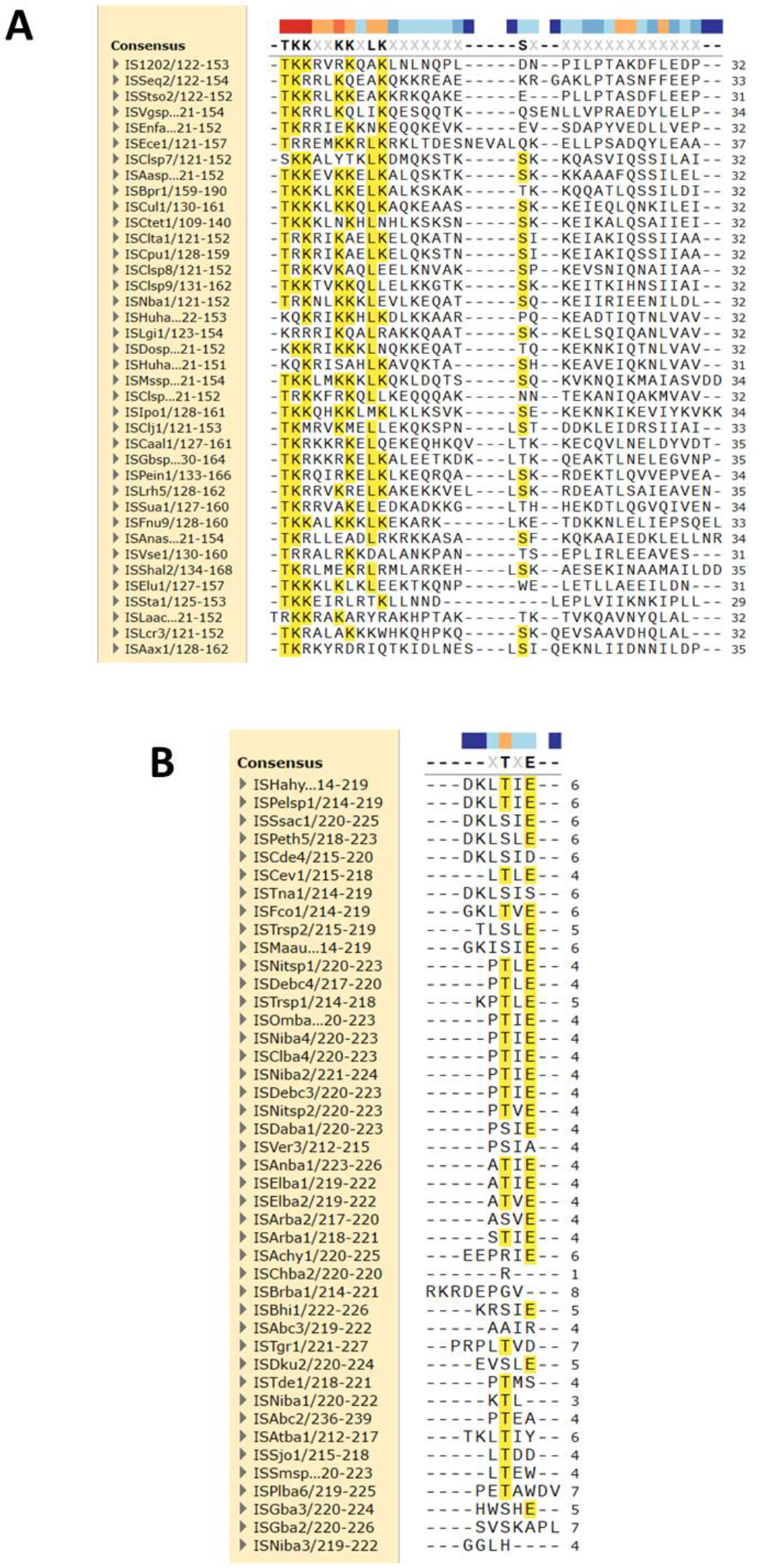
Alignment of indels sequences. The alignments were prepared using SnapGene. The color code above indicates the degree of conservation. A : IS*1202* group indel. B : IS*Tde1* group indel.

**Supplementary Table 1. A Summary of the entire IS*1202* family**.

