## Supplementary figures and images for "A subclass of the IS*1202* family of bacterial insertion sequences targets XerCD recombination sites"

### Supplementary Figure 1

#### Supplementary Figure 1

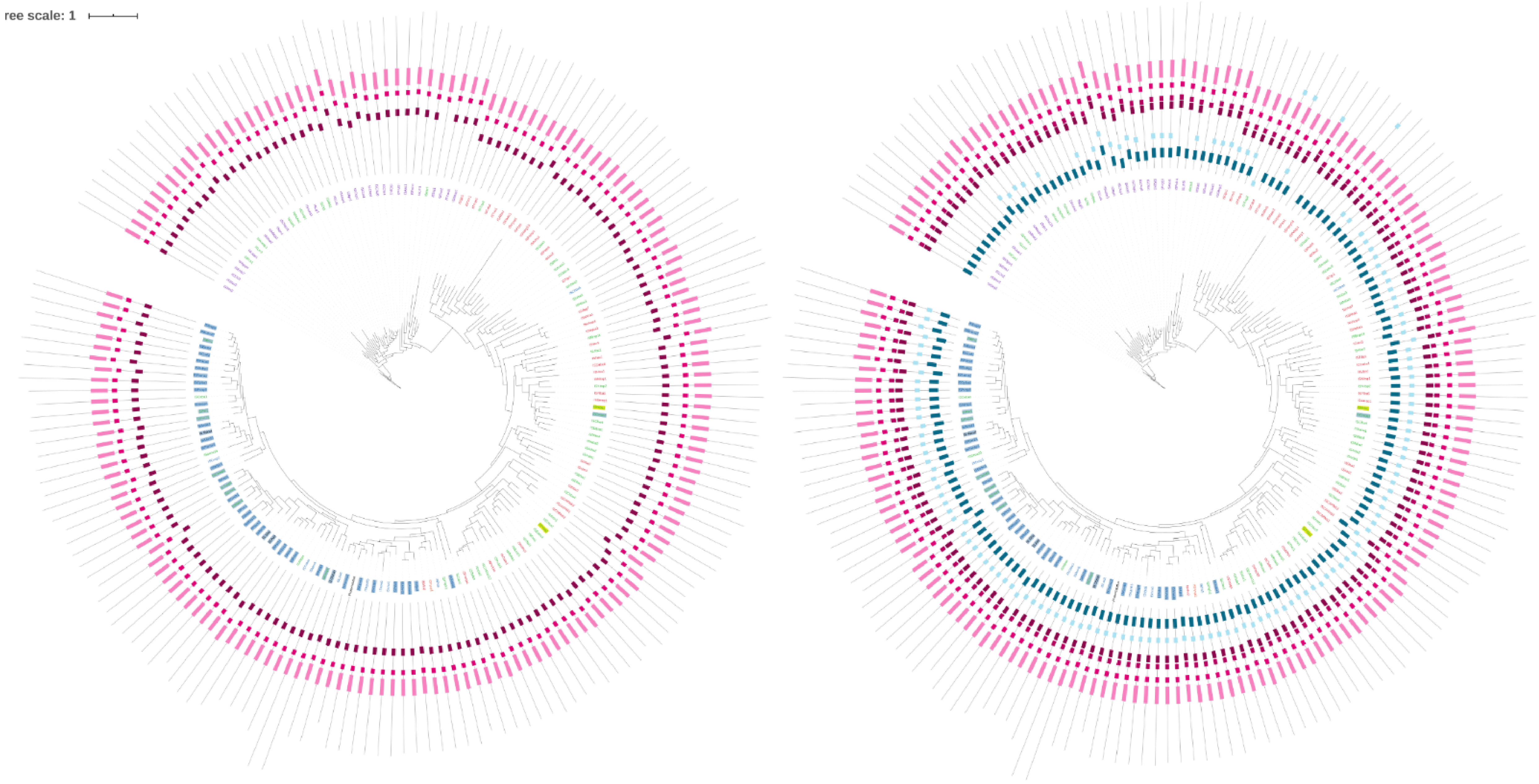

### Supplementary Figure 2

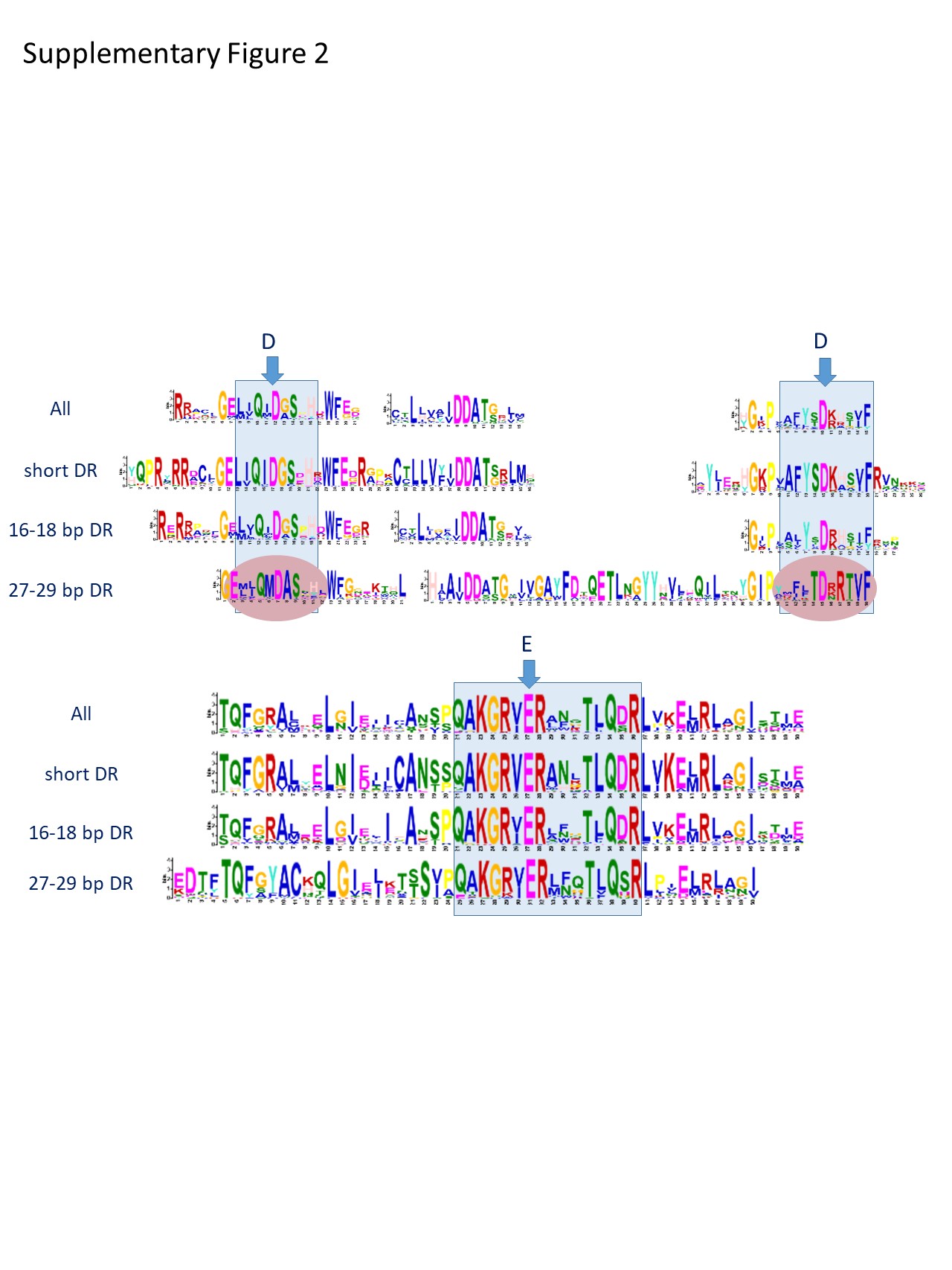

### Supplementary Figure 3

#### Supplementary Figure 3

A

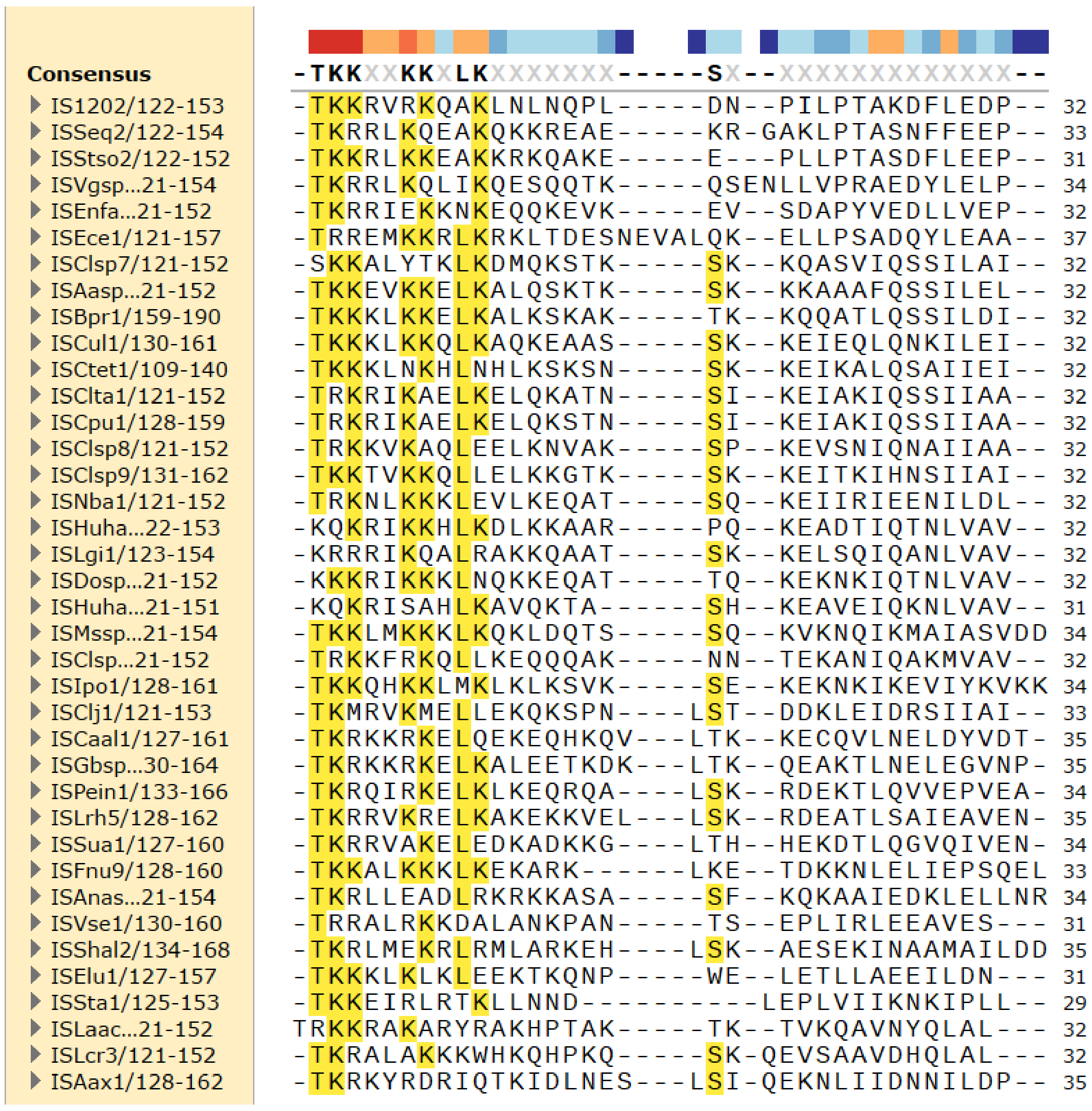


B

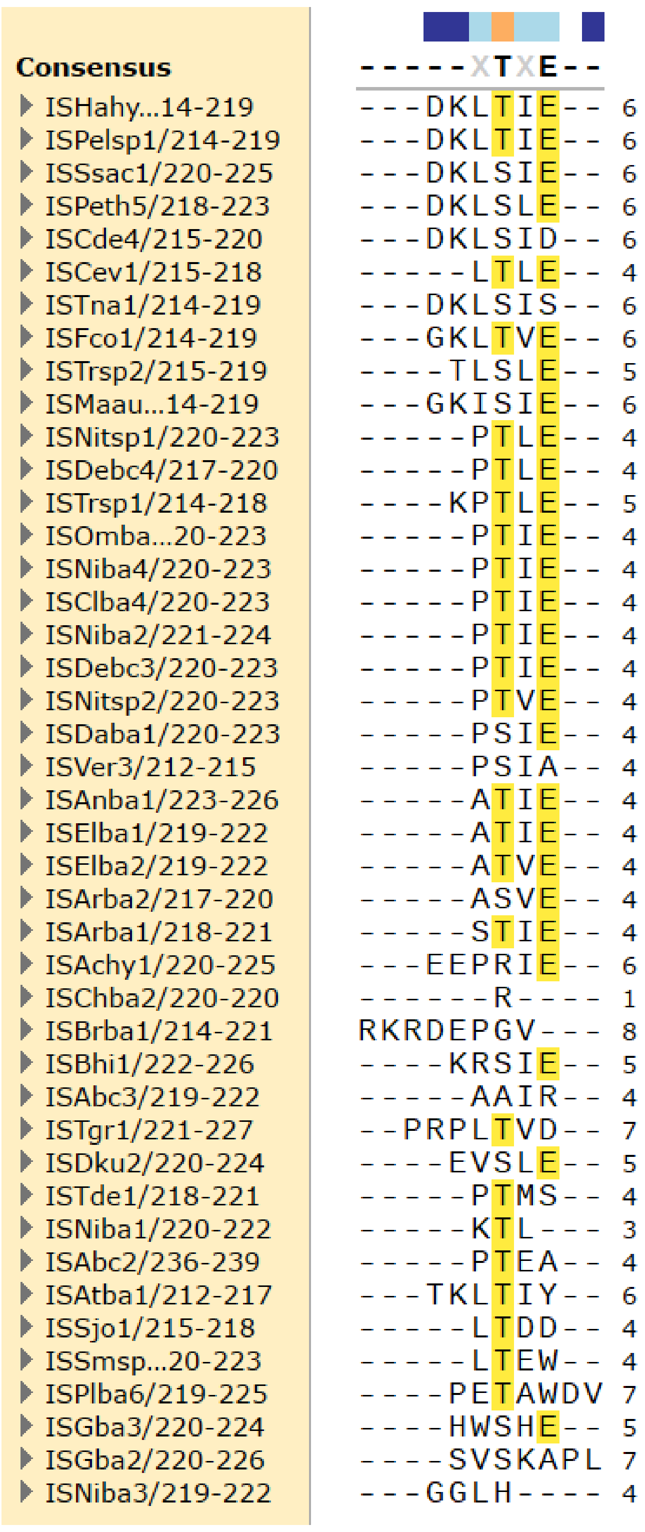
